# *NRMLncR,* a myocyte-enriched long non-coding RNA, enhances myogenesis in mouse

**DOI:** 10.64898/2026.02.11.704964

**Authors:** Yufen Li, Yumei Zhou, Qing Ying Li, Justine Arrington, Mostafa F. Abdelhai, Yubo Wang, Junxiao Ren, Yung-Yu Cheng, Mingyi Xie, W. Andy Tao, Shihuan Kuang, Feng Yue

**Affiliations:** Department of Animal Sciences, University of Florida, Gainesville, FL 32611, USA; Department of Biochemistry, Purdue University, West Lafayette, IN 47907, USA; Mass Spectrometry Core, Salk Institute for Biological Studies, La Jolla, CA 92037, USA; Department of Biochemistry and Molecular Biology, University of Florida, Gainesville, FL 32610, USA; UF Genetics Institute, University of Florida, Gainesville, FL 32611, USA; Department of Animal Sciences, Purdue University, West Lafayette, IN 47907, USA; Department of Orthopaedic Surgery, Duke University, Durham, NC 27710, USA; UF Myology Institute, University of Florida, Gainesville, FL 32611, USA

**Keywords:** Myoblast, myogenesis, long non-coding RNA, myogenic regulatory factor, Notch signaling, CELF1

## Abstract

Myogenesis is a stepwise process encompassing myogenic progenitor proliferation, lineage commitment, differentiation, myocyte fusion, and myotube maturation, and it is orchestrated by myogenic regulatory factors (MRFs) together with signaling pathways that coordinate these transitions. Long noncoding RNAs (lncRNAs) have emerged as important regulators of muscle development and regeneration, yet how lncRNAs integrate with canonical signaling networks to shape myogenic progression remains incompletely defined. Here, we identify a novel myocyte-enriched, Notch-repressed myogenic lncRNA (*NRMLncR,* known as *A930003A15Rik*), as a previously uncharacterized regulator of mouse myogenesis. The expression of *NRMLncR* is robustly induced during primary myoblast activation and differentiation. Loss-of-function analyses show that knockdown of *NRMLncR* impairs myogenic differentiation, accompanied by reduced expression of key myogenic genes. In contrast, adenovirus-mediated overexpression of *NRMLncR* enhances myogenic differentiation *in vitro* and is associated with increased muscle fiber size *in vivo*. Mechanistically, MyoD and MyoG occupy the *NRMLncR* promoter and promote its transcription during myogenic differentiation. *NRMLncR* knockdown alerts the transcription of nearby genes, suggesting its function through a cis-regulatory mechanism. RNA pull-down assays further identify an interaction between *NRMLncR* and the RNA-binding protein CELF1. Together, these findings establish *NRMLncR* as a novel Notch-associated lncRNA that promotes myogenic differentiation and provide insight into lncRNA-dependent regulation of the myogenic program.

## 1 | INTRODUCTION

Skeletal muscle regeneration relies on efficient myogenesis driven by the activation and coordinated fate decisions of resident muscle satellite cells (SCs).^1–3^ In uninjured muscle, SCs are maintained in a quiescent state. Upon injury, SCs rapidly activate, re-enter the cell cycle, and expand to generate proliferating muscle progenitor cells or myoblasts that fuel repair.^4–6^ During regeneration, a large fraction of activated myoblasts committed to myogenic differentiation as myocyte and fuse to form or repair multinucleated myofibers, whereas a distinct subset undergoes self-renewal and returns to quiescence to preserve the stem cell pool.^4–6^ This stepwise myogenic program is orchestrated by myogenic regulatory factors (MRFs), with MyoD promoting lineage commitment, myogenin (MyoG) driving terminal differentiation, and MRF4 supporting myofiber maturation and stabilization of the muscle architecture.^7–11^ Maintaining an appropriate balance between SC self-renewal and differentiation is therefore essential for sustained regenerative capacity across repeated cycles of injury over lifetime.^12–14^ Multiple extrinsic and intrinsic regulators, including Notch,^15–20^ Wnt,^21–24^ p38/MAPK,^25–27^ AMPK,^28–31^ and PI3K/AKT signaling pathways,^32–35^ integrate environmental cues with the core myogenic transcriptional network to guide these transitions.^6, 14^ Among these pathways, Notch signaling is a central determinant of SC quiescence and is required to maintain self-renewal and prevent premature differentiation.^18–20^ Accordingly, impaired Notch signaling is associated with defective muscle regeneration and pathology.^36^ However, despite the recognized importance of Notch in SC fate control, the downstream regulatory mechanisms that connect Notch activity to stage-specific myogenic progression remain incompletely understood.

Long non-coding RNAs (lncRNAs) have emerged as important regulatory components of skeletal myogenesis, contributing to the coordinated progression from myoblast expansion to differentiation, myofiber formation, and regeneration.^37, 38^ Increasing evidence indicates that many lncRNAs exhibit stage-specific expression patterns during myogenic progression and that perturbation of individual lncRNAs can significantly impact muscle development and repair.^37, 38^ Among the best-characterized examples, *linc-MD1* and *lnc-mg* control the timing of muscle differentiation by functioning as a competing endogenous RNA to fine-tune pro-differentiation gene expression programs.^39, 40^ Additional lncRNAs regulate myogenesis by influencing chromatin state and transcriptional output at myogenic loci, exemplified by *Linc-RAM* which cooperates with MyoD to support activation of myogenic genes,^41^ *lncMREF* which promotes differentiation and regeneration through interaction with the Smarca5/p300 complex,^42^ and *Linc-YY1* which modulates myogenic differentiation and regeneration through functional interaction with YY1.^43^ Epigenetic control of myogenesis by lncRNAs is further highlighted by lncRNA *Dum*, which supports myogenic differentiation and regeneration by coordinating DNA methylation-dependent regulation of myogenic programs.^44^ In contrast, lncRNA *SYISL* functions as a repressor of myogenesis by interacting with polycomb repressive complex 2 (PRC2), and its genetic deletion in mice increases muscle fiber density and muscle mass.^45^ In parallel, lncRNAs also impact the expansion of the myogenic progenitor pool, as illustrated by *SAM*, which promotes myoblast proliferation and is required for efficient regenerative responses *in vivo*.^46^ Collectively, these studies establish lncRNAs as integral regulators of myogenic progression, yet the functions of most muscle-associated lncRNAs remain incompletely defined, underscoring the need to further delineate lncRNA-dependent regulatory networks in myogenesis and regeneration.^37, 38^

In this study, we sought to identify Notch-repressed lncRNAs that contribute to the regulation of myogenic differentiation. Using microarray profiling of Notch-activated myoblasts, we identified *NRMLncR* (*A930003A15Rik*) as a candidate lncRNA that is inversely associated with Notch activity and dynamically regulated during myogenesis and muscle remodeling contexts. We then characterized *NRMLncR* with respect to its coding potential, transcriptional regulation by the myogenic program, and functional contribution to differentiation and muscle growth. Finally, we explored potential molecular mechanisms by identifying *NRMLncR*-interacting proteins, revealing an association with the RNA-binding factor CELF1. These findings establish *NRMLncR* as a Notch-associated lncRNA regulator and expand the repertoire of noncoding transcripts linked to myogenic control.

## 2 | MATERIALS AND METHODS

### 2.1 | Animals

All procedures involving animals were conducted in compliance with National Institutes of Health and Institutional guidelines with approval by the Purdue Animal Care and Use Committee. *Rosa26-N1ICD* (stock# 006850), *mdx* (stock# 001801) and *C57BL/6J* (stock# 000664) mice were purchased from the Jackson Lab. *Tg: Pax7-nGFP* mouse was provided by Dr. Shahragim Tajbakhsh (Institut Pasteur).^47^ Genotyping of the mouse was performed by genomic DNA isolated from ear and were screened by polymerase chain reaction (PCR) using primers and program following the protocols provided by the supplier. Mice were housed under standard laboratory conditions with free access to food and water. If not stated differently, 2- to 6-month-old adult mice were used for all experiments. Both male and female mice were used for each specific experiment.

### 2.2 | Muscle injury and regeneration

Muscle injury was induced by cardiotoxin (CTX) injection. Briefly, adult mice were anesthetized using ketamine-xylazine, and 50 μL of 10 μM CTX (Sigma-Aldrich, cat#217503) was injected into the tibialis anterior (TA) muscle. Injured muscles were harvested at the indicated time points for subsequent analyses of muscle regeneration.

### 2.3 | Fluorescence-activated cell sorting (FACS)

*Tg: Pax7-nGFP* reporter mice were used for the isolation of muscle SCs. Muscle injury was induced by CTX injection on day 3 to obtain activated SCs, while quiescent SCs were isolated from the contralateral uninjured muscle. Hindlimb muscles were harvested, and single-cell suspensions were prepared as previously described.^48^ Cells were stained with a Zombie Violet Live/Dead dye (BioLegend, cat#423113) to exclude dead cells and subjected to FACS analysis. SCs were sorted based on GFP fluorescence and Violet-negative signals. Sorted cells were examined under a Leica DMi6000B microscope to confirm cell purity.

### 2.4 | Cell culture and differentiation

Primary myoblasts were isolated from hindlimb muscles of 2-4-week-old wild-type mice as previously described.^49^ Briefly, muscles were dissected, minced, and digested in a mixture of type I collagenase (Worthington, cat#LS004197) and Dispase II (Roche, cat#4942078001). Digestion was stopped by adding Ham’s F-10 medium (Gibco, cat#11550043) supplemented with 20% fetal bovine serum (FBS, Corning, cat#35-011-CV). The cell suspension was filtered to remove debris, centrifuged, and plated on collagen-coated dishes. Cells were maintained at 37 °C in a humidified incubator with 5% CO₂ in growth medium consisting of Ham’s F-10 medium supplemented with 20% FBS, 4 ng mL^-1^ basic fibroblast growth factor (bFGF, Gibco, cat#PHG0263), and 1% penicillin–streptomycin (P/S, Gibco, cat#15-140-122). For differentiation, primary myoblasts were seeded on BD Matrigel (Corning, cat#354234)-coated plates and cultured in differentiation medium containing DMEM (Gibco, cat#11995065) supplemented with 2% horse serum (HyClone, cat#SH30074) and 1% P/S.

C2C12 cells were maintained at 37 °C in a humidified incubator with 5% CO₂. C2C12 cells were cultured in growth medium consisting of DMEM supplemented with 10% FBS and 1% P/S. For myogenic differentiation, C2C12 cells were switched to differentiation medium (DMEM supplemented with 2% horse serum and 1% P/S) at the indicated time points and maintained for the duration specified in each experiment. HEK293T/HEK293A cells were cultured in DMEM supplemented with 10% FBS and 1% P/S and were used for plasmid transfection and luciferase assays as indicated.

### 2.5 | Lentivirus-mediated gene stable knockdown

Lentiviral short hairpin RNAs (shRNAs) targeting *NRMLncR* were designed using an online RNAi design tool (Thermo Fisher Scientific) as shown in Supplemental Table 1. The shRNA sequences were cloned into the pLKO.1 vector (Addgene, cat#8453). Lentiviral particles were produced by co-transfecting HEK293T cells with the pLKO.1-shRNA plasmid targeting *NRMLncR* or pLKO.1-Scramble (a gift from David Sabatini, Addgene, cat#1864), together with the packaging plasmids pCMV-VSV-G (Addgene, cat#8454) and pCMV-ΔR8.2 (Addgene, cat#8455). Viral supernatants were collected 48 h after transfection, filtered through a 0.45-μm membrane, and used to infect C2C12 cells and SC-derived primary myoblasts at approximately 30% confluence. After 48 h, infected cells were selected with puromycin (1 μg mL^-1^, Fisher Scientific, cat#BP2956100) for two passages to establish stable knockdown cell lines. Knockdown efficiency was confirmed by quantitative RT-PCR as indicated.

### 2.6 | Adenovirus-mediated gene overexpression

Adenovirus-Cre (Ad-Cre) and adenovirus-GFP (Ad-GFP) were gifts from Dr. Yong-Xu Wang (University of Massachusetts Medical School, MA, USA). Recombinant adenoviruses expressing *NRMLnc* were generated using the AdEasy system according to protocols.^50, 51^ The *NRMLnc* cDNA was amplified from mouse primary myoblasts and cloned into the pAdTrack-CMV vector. The formed pAdTrack-*NRMLnc* (pAdTrack-CMV as the control) plasmid were digested by PmeI (NEB, cat#R0560S), and then transfected into BJ5183-AD1 competent cells (Agilent, Cat#200157) with pAdEasy-1 plasmid. The positive recombinant plasmid was detected by PacI (NEB, cat#R0547S) digestion. For adenovirus generation, HEK293A cells (60-70% confluent) in 10-cm culture dishes were transfected with 4 μg of PacI-digested recombinant plasmid using Lipofectamine 2000 (Life Technologies, cat#11668019). After 10 days of transfection, the cells were collected and recombinant adenovirus was released by freeze-thaw method. To increase the titers of adenovirus, two more rounds of infection were adapted to amplify the recombinant virus, and the titers were then determined by the expression of GFP. For *in vitro* experiments, SC-derived primary myoblasts were infected with adenovirus expressing Ad-NRMLnc or control Ad-GFP for 12 hours and cultured with growth and differentiation medium for following studies.

### 2.7 | Adenovirus purification and intramuscular administration

Recombinant adenoviruses were used for *in vivo* gene delivery to skeletal muscle as previously described.^50^ Adenoviruses were amplified in HEK293A cells from thirty 10-cm culture dishes through stepwise scale-up infection and harvested when ∼50% of infected cells detached. Cells were collected and pelleted by centrifugation, and viruses were released by repeated freeze-thaw-vortex cycles to generate clarified viral lysates. For high-titer purification, cleared lysates were mixed with CsCl and subjected to ultracentrifugation using a Beckman SW 41 Ti rotor for 18 h at 176,000g to separate viral particles, and the visible virus band was collected using a syringe. Purified virus was then mixed with adenovirus storage buffer (10 mM Tris-HCl, pH 8.0, 100 mM NaCl, 0.1% BSA, 50% glycerol), filter-sterilized, and stored at −80 °C until use.

Purified adenoviral particles were diluted in sterile saline and administered by intramuscular injection into the TA muscle of mice. In brief, mice were anesthetized by intraperitoneal injection of a ketamine/xylazine cocktail. The hindlimb was shaved and disinfected with 70% ethanol, and 20 μL of purified Ad-NRMLnc virus (1 x10^11^ particles) was injected into the TA muscle using a sterile 0.5-cc disposable insulin syringe. Control animals received an equivalent dose and volume of control Ad-GFP virus using the same procedure. Following injection, mice were placed on a warmed pad until fully recovered and then returned to the animal facility, with injected animals housed separately from non-injected animals. Tissues were collected at the indicated time points for downstream analyses.

### 2.8 | Total RNA extraction and real-time PCR

Total RNA was extracted from cultured cells or mouse tissues using TRIzol reagent (Thermo Fisher Scientific, cat#15596018) according to the manufacturer’s instructions. RNA concentration and purity were assessed spectrophotometrically. Complementary DNA (cDNA) was synthesized from total RNA using random primers with M-MLV reverse transcriptase (Invitrogen, cat#28-025-021). Quantitative real-time PCR (qRT-PCR) was performed using SYBR Green Master Mix (Roche, cat#4913850001) on a LightCycler 96 Real-Time PCR System (Roche). Relative gene expression levels were calculated using the 2^⁻ΔΔCt^ method and normalized to the indicated housekeeping genes.^52^ Primer sequences are listed in Supplemental Table 1.

### 2.9 | Microarray analysis

Ad-Cre and Ad-GFP adenovirus were infected to 10-cm plates of Rosa-NICD primary myoblasts. After 6 h, fresh Ham’s F-10 growth medium was added to replace the virus cultures and cells were collected after an additional 48 h. Microarray was performed as described in published protocol.^20, 53^ RNA was extracted from cultured primary myoblasts infected with Ad-GFP and Ad-Cre adenovirus. Gene expression was analyzed by microarray with Agilent SurePrint G3 Mouse GE 8 × 60 K chip (Agilent, cat# G4858A). The list of significantly changed genes with a fold change ≥1.5-fold was used for Gene Ontology (GO) and Kyoto Encyclopedia of Genes and Genomes (KEGG) pathway analysis.

### 2.10 | Chromatin immunoprecipitation (ChIP)-qPCR

ChIP-qPCR was performed to examine the binding of MRFs to the *NRMLncR* promoter in differentiated myoblasts. Briefly, cells were crosslinked, lysed, and sonicated to generate fragmented chromatin. Chromatin was immunoprecipitated using antibodies against MyoD (Santa Cruz Biotechnology, cat#sc-377460) and MyoG (DSHB, cat#F5D), with mouse IgG (Santa Cruz Biotechnology, cat#sc-2025) serving as a negative control. Immune complexes were captured using protein A/G agarose beads (Santa Cruz Biotechnology, cat#sc-2003) and washed extensively. Crosslinks were reversed by proteinase K (Roche, cat#3115836001) treatment, and DNA was purified using phenol-chloroform extraction. Enriched DNA fragments were analyzed by quantitative PCR using primers (Supplemental Table 1) targeting the E-box regions of the *NRMLncR* promoter.

### 2.11 | Protein extraction and western blot analysis

Protein samples were extracted from cultured cells or mouse tissues using RIPA lysis buffer supplemented with protease inhibitors (Sigma-Aldrich, cat#P8340). Protein concentrations were determined using Pierce BCA Protein Assay Reagent (Pierce Biotechnology, cat#PI23225). Equal amounts of protein were mixed with Laemmli sample buffer (Bio-Rad, cat#1610747), boiled for 10 min, and separated by SDS-PAGE. Proteins were transferred onto PVDF membranes (Millipore, cat#ISEQ00010). Membranes were blocked in 5% non-fat milk and incubated with primary antibodies at 4 °C overnight, followed by incubation with appropriate secondary antibodies at room temperature for 1 h. The following primary and secondary antibodies were used: MyoG (DSHB, cat#F5D), MF20 (DSHB, cat#MF-20), FLAG (Sigma-Aldrich, cat#F1804), GAPDH (Santa Cruz Biotechnology, cat#sc-32233), HRP AffiniPure goat anti-mouse IgG (Jackson ImmunoResearch, cat#115-035-003). Immunodetection was performed using enhanced chemiluminescence western blotting substrate (Santa Cruz Biotechnology, cat#sc-2048) and detected with a FluorChem R System (Proteinsimple).

### 2.12 | Immunofluorescence staining

Cells were fixed with 4% paraformaldehyde for 10 min at room temperature and quenched with 100 mM glycine for 10 min. After washing with PBS, cells were blocked in blocking buffer for 1 h at room temperature. Samples were incubated with primary antibodies at 4 °C overnight, followed by incubation with appropriate fluorescent secondary antibodies and DAPI at room temperature for 1 h. The following primary and secondary antibodies were used: anti-MyoG (DSHB, cat#F5D), MF20 (DSHB, cat#MF-20), FLAG (Sigma-Aldrich, cat#F1804), Alexa 568 goat anti-mouse IgG1 (Invitrogen, cat#A-21124), and Alexa 647 goat anti-mouse IgG2b (Invitrogen, cat#A-21242). Images were acquired with a Leica DM6000 microscope with a 10× or 20× objective, and image analysis was performed using ImageJ.

### 2.13 | Dual-Luciferase assay

pGL3-NRMLncR vector was constructed by cloning *NRMLncR* promoter (containing 1,595 bp upstream from transcription start site, TSS) into pGL3-basic vector (Addgene, cat#212936). HEK293 cells were seeded in 48-well plates and co-transfected with the pGL3-NRMLncR promoter luciferase reporter plasmid (or pGL3-basic-Empty as control), expression vectors for pRS2-RBPJκ-VP16, pRS2-RBPJκ-DBM (a gift from Dr. Mark Mercola),^54^ pcDNA-MyoD, pcDNA-MyoG, or pcDNA-MRF4 (or pcDNA3-Empty as control), and a Renilla luciferase plasmid as an internal control. After 48 h, cells were harvested, and luciferase activities were measured using the Dual-Luciferase Reporter Assay System (Promega, cat#E1960) following the manufacturer’s instructions. Firefly luciferase activity was normalized to Renilla luciferase activity to calculate the relative reporter activity.

### 2.14 | RNA pull-down assay

RNA pull-down assays were performed following previous reports^55^ to identify proteins associated with sense or antisense *NRMLncR*. Briefly, sense and antisense *NRMLncR* DNA was cloned into pGEM-T vector (Promega, cat#A3600) with primers shown in Supplemental Table 1, linearized and purified. Biotinylated sense and antisense *NRMLncR* RNAs were synthesized using the MegaScript T7 Transcription Kit Plus (Invitrogen, cat#A57622-25) with linearized pGEM-T-NRMLncR-sense and pGEM-T-NRMLncR-antisense as template. The biotin-16-UTP (Roche, cat#86303-26-6) was used to label *NRMLncR* RNA with the ratio of 1:3 to UTP. Biotinylated RNAs were incubated with C2C12 myotube lysates under rotation at 4 °C to allow formation of RNA-protein complexes. Complexes were captured using streptavidin-coated beads and washed extensively to reduce non-specific binding. Bound proteins were eluted and analyzed by SDS-PAGE followed by Coomassie blue staining and/or immunoblotting. For unbiased identification of candidate binding proteins, eluates were subjected to mass spectrometry as described previously.^56^, and data was analyzed using MaxQuant v1 software.

### 2.15 | Statistical analysis

All quantitative data are presented as mean ± standard error of the mean (SEM) from at least three independent biological replicates except two replicates for microarray analysis. Statistical significance between two groups was determined using a two-tailed Student’s *t*-test, One-way ANOVA or Two-way ANOVA with Tukey’s, Kruskal-Wallis or Šídák multiple comparisons test. A *P* value < 0.05 was considered statistically significant.

## 3 | RESULTS

### 3.1 | Identification of a novel LncRNA negatively associated with Notch involved in myogenic differentiation

To identify downstream genes regulated by Notch signaling during myogenesis, we isolated primary myoblasts from *Rosa-NICD* mice and infected them with Ad-Cre to induce overexpression (OE) of the Notch1 intracellular domain (NICD), using Ad-GFP-infected myoblasts as controls (Fig. 1A). Microarray profiling revealed 1,491 differentially expressed protein-coding genes in Notch OE myoblasts compared with controls, including 819 upregulated and 672 downregulated genes (Fig. 1B, Supplemental Datasheet 1). Gene Ontology (GO) analysis showed that upregulated genes were enriched for biological processes associated with extracellular matrix organization, positive regulation of cell migration, cell adhesion, response to hypoxia, glutathione metabolic process, and the Notch signaling pathway (Fig. 1C). In contrast, downregulated genes were strongly associated with myogenic programs, including skeletal muscle cell differentiation, muscle contraction, chromatin remodeling, myoblast fusion, cell fate commitment, and biological processed related to membrane potential regulation and intracellular calcium ion homeostasis (Fig. 1C). Consistently, KEGG pathway analysis of upregulated genes identified significant enrichment of focal adhesion, ECM-receptor interaction, glutathione metabolism, Notch signaling, and HIF-1 signaling pathways (Fig. 1D). Together, these transcriptomic changes are consistent with the established role of Notch signaling in maintaining myogenic progenitors, promoting self-renewal, and suppressing premature differentiation.

**Figure 1.**
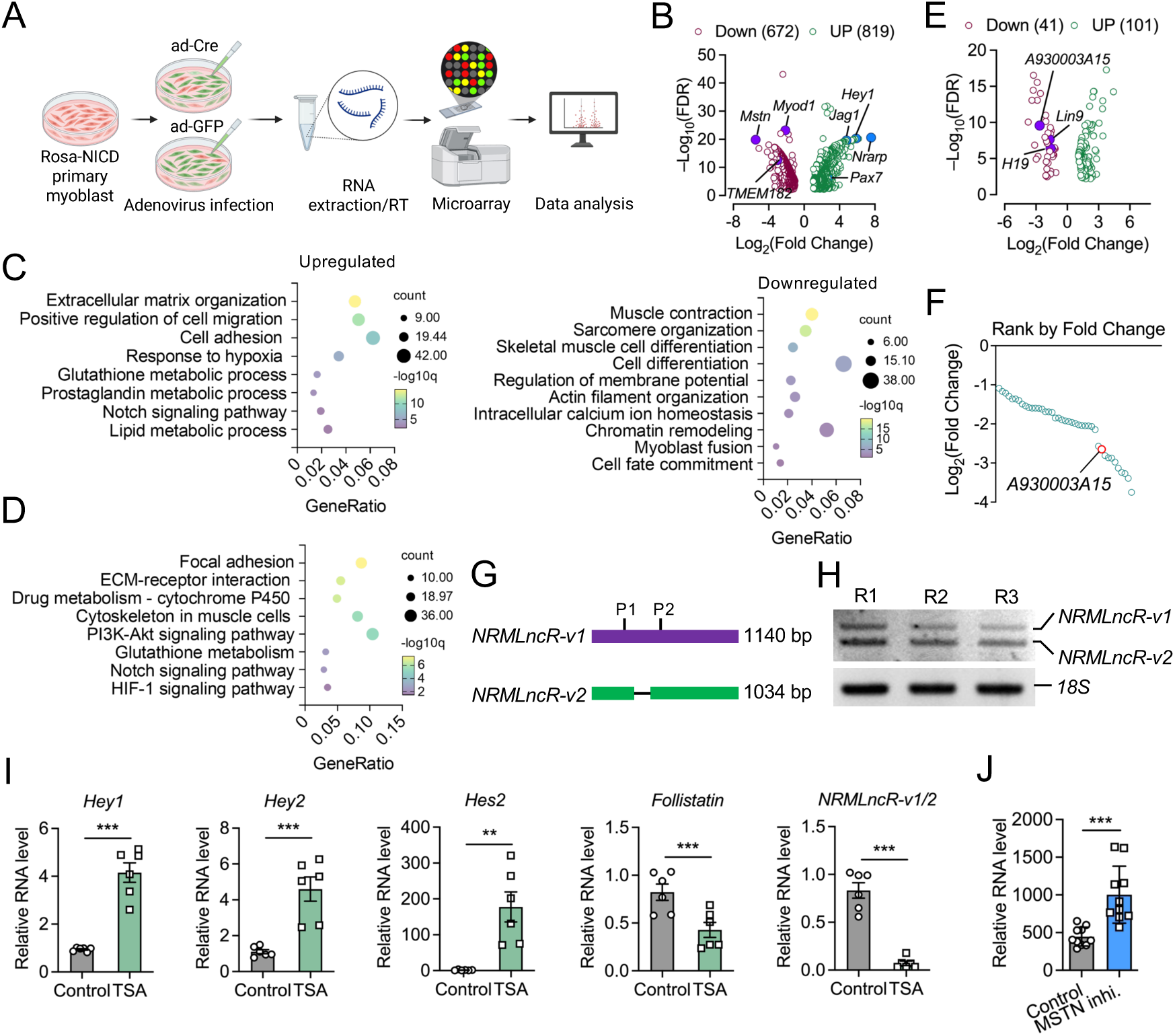
Identification of a Notch-responsive myogenic lncRNA *NRMLncR* in myoblasts. (A) Schematic illustration of experimental design for microarray analysis of Rosa-NICD primary myoblasts infected with Ad-GFP (control) or Ad-Cre to activate Notch signaling. (B) Volcano plot showing differentially expressed genes between Ad-Cre- and Ad-GFP-treated primary myoblasts. (C and D) Dot plots showing enrichment of biological process pathways from GO (C) and KEGG (D) analyses. (E) Differentially expressed lncRNAs identified from microarray profiling of Notch-activated primary myoblasts. (F) Rank of downregulated lncRNAs in Notch-activated primary myoblasts based on the fold changes. (G) Schematic representation of the two NCBI annotated *NRMLncR* transcripts, *NRMLncR-v1* and *NRMLncR-v2*. (H) Electrophoresis analysis showing *NRMLncR-v1* and *NRMLncR-v2* expression in primary myoblasts. Lanes indicates three replicates. (I) Expression of Notch-target genes and total *NRMLncR-v1/2* in C2C12 cells after pharmacological activation of Notch signaling by trichostatin A (TSA). n=5. (J) Expression of total *NRMLncR-v1/2* following MSTN inhibition. Dataset: GSE13707. n=10. Data represent mean ± SEM. Student’s *t*-test; \*\**p* < .01; and \*\*\**p* < .001.

Among the differentially expressed transcripts, we identified 142 annotated lncRNAs with differentially expression in Notch OE myoblasts (Fig. 1E, Supplemental Datasheet 1). Of these, 41 were downregulated and 101 upregulated (Fig. 1E). While most of the lncRNAs were uncharacterized, the well-studied lncRNAs H19 was significantly downregulated in Notch OE myoblasts, consistent with its known role in promoting myogenic differentiation^57,58^. Notably, the gene *A930003A15Rik* ranked as the top 10 downregulated lncRNA in Notch OE myoblasts (log2FC = –2.65) (Fig. 1E, F). We therefore named *A930003A15Rik* Notch-repressed myogenic LncRNA (*NRMLncR*). According to the NCBI database, *A930003A15Rik* has two annotated transcript variants, *NRMLncR-v1* (1,140 bp) and *NRMLncR-v2* (1,034 bp), with *NRMLncR-v2* lacking 106 nucleotides (Fig. 1G). Both variants were cloned from primary myoblasts and detected by gel electrophoresis (Fig. 1H). To validate whether *NRMLncR* is responsive to Notch activation, we treated C2C12 cells with trichostatin A (TSA), a pharmacological inhibitor of pan-HDAC activating Notch signaling.^59^ TSA treatment significantly upregulated the Notch downstream genes *Hey1*, *Hey2*, and *Hes2*, while downregulating the *Follistatin* mRNA expression (Fig. 1I). Strikingly, the expression of total *NRMLncR-v1*/2 remarkably decreased 11.7-fold in TSA-treated C2C12 cells (0.832 in control versus 0.071 in TSA-treated cells) (Fig. 1I). Conversely, inhibition of myostatin (MSTN, GSE13707) increased *NRMLncR* expression (Fig. 1J). Together, these results indicate that *NRMLncR* expression is responsive to Notch pathway activity.

### 3.2 | Dynamic expression of *NRMLncR* correlates with the myogenic program

To characterize the expression pattern of *NRMLncR*, we examined its levels across multiple tissues, disease models, and postnatal stages in mice. Tissue profiling from a public RNA-seq dataset (GSE9954) showed that *NRMLncR-v1/2* was predominantly expressed in the mouse eye, with high expression in the thymus and moderate expression in the diaphragm and skeletal muscle, compared with other organs (Fig. 2A). In dystrophic *mdx* mice, expression of *NRMLncR-v1/2* and *NRMLncR-v1* in skeletal muscle was significantly higher than in wild-type (WT) controls, with 4.1- and 4.5-fold increases, respectively (Fig. 2B). In addition, both *NRMLncR-v1/2* and *NRMLncR-v1* were highly expressed in neonatal skeletal muscle and gradually decreased with aging, a pattern consistent with the expression of *Pax7* and *MyoG* during neonatal myogenesis (Fig. 2C). These data suggest that *NRMLncR* expression is associated with myogenic program rather than myofiber maturation.

**Figure 2.**
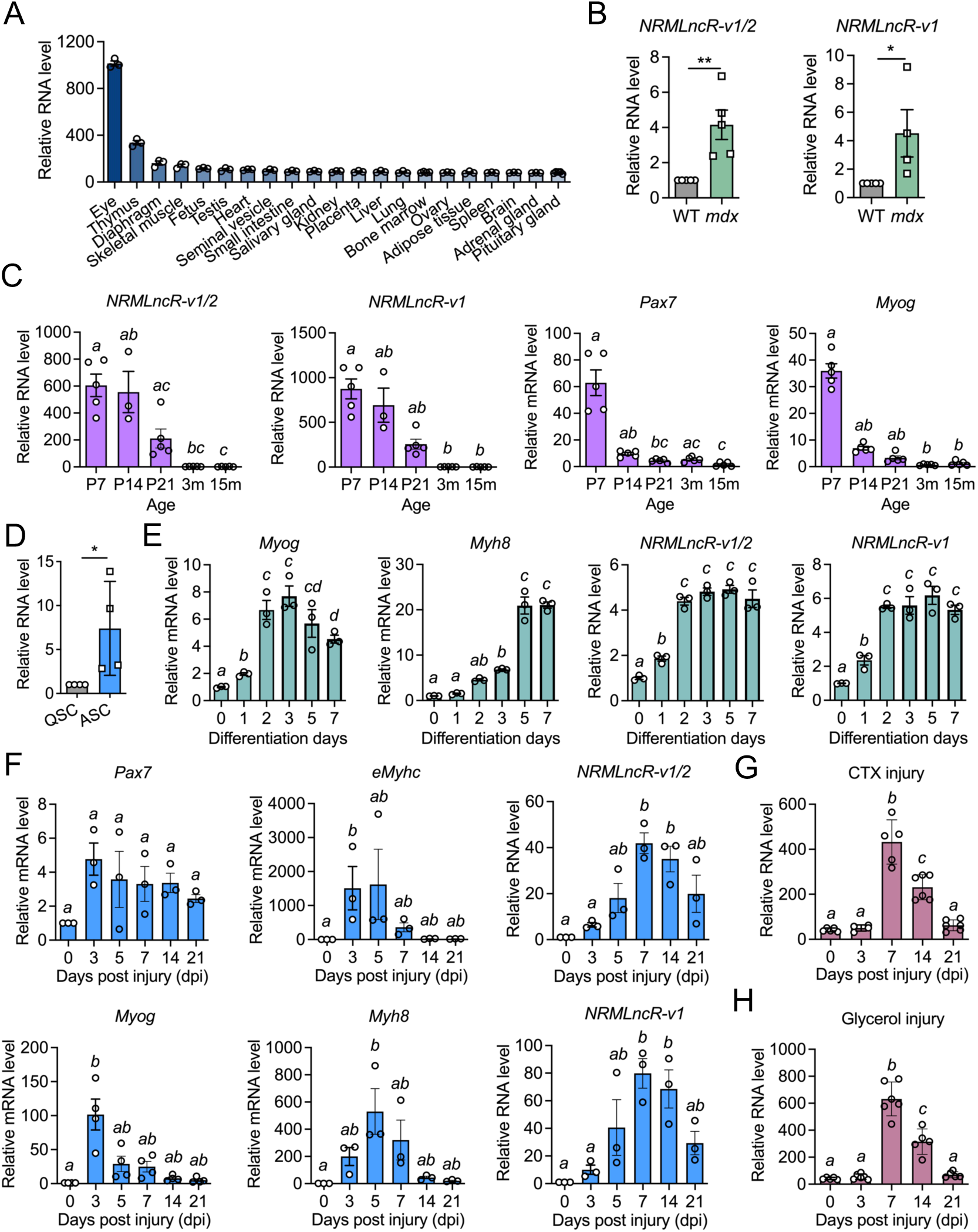
Dynamic expression of *NRMLncR* corelates with myogenic differentiation. (A) Expression levels of *NRMLncR-v1/2* across different mouse tissues. Dataset: GSE9954. n=3. (B) Relative mRNA abundance of *NRMLnc-v1/2* in skeletal muscle from wild-type (WT) and *mdx* mice. *NRMLncR-v1/2*: n=5; *NRMLncR-v1*: n=5 for WT, n=4 for *mdx*. (C) Expression of *NRMLncR-v1/2*, *NRMLncR-v1*, *Pax7*, and *Myog* in skeletal muscle from postnatal day 7 to 15 months of age. n=3-5. (D) Expression of *NRMLncR-v1/2* in quiescent and activated satellite cells (SCs). n=4. (E) Expression of *NRMLncR-v1/2*, *NRMLnc-v1*, and myogenic differentiation markers (*Myog* and *Myh8*) in C2C12 cells after 7 days of differentiation. n=3. (F) Expression of *NRMLncR-v1/2*, *NRMLncR-v1*, *Pax7*, *eMyhc*, *Myog*, and *Myh8* following cardiotoxin (CTX)-induced muscle injury in mice. n=3. (G and H) *NRMLncR* expression profiles in an independent CTX- (G) and glycerol-induced (H) muscle injury model. Dataset: GSE45577. n=5-6. Data represent mean ± SEM. Student’s *t*-test (B, D): \*\**p* < .01; and \*\*\**p* < .001. One-way ANOVA multiple comparisons test (C, E-H): groups with different letters are statistically different (*p*<.05).

To further test whether *NRMLncR* expression correlates with the myogenic program, we examined its expression during SC activation, myogenic differentiation, and muscle regeneration. Notably, *NRMLncR* expression was increased 7.4-fold in activated SCs FACS-sorted from CTX-injured *Tg: Pax7-nGFP* mice compared with quiescent SCs isolated from uninjured muscles (Fig. 2D). During *in vitro* myogenic differentiation of C2C12 myoblasts, the myogenic differentiation marker *Myog* reached peak expression at 2 and 3 days after differentiation and progressively decreased at 5 and 7 days, whereas the myotube marker *Myh8* gradually increased from days 0-3, showed a sharp increase at 5 days, and remained at high levels at 7 days (Fig. 2E). In parallel, the expression of both *NRMLncR-v1/2* and *NRMLncR-v1* increased beginning at day 1 of differentiation, reached higher levels by day 2, and remained stable thereafter (Fig. 2E). These results suggest a potential role for *NRMLncR* in both early and late myogenesis.

Following CTX-induced muscle injury, the myogenic marker genes exhibited dynamic expression patterns, with *Pax7* and *Myog* peaking at 3 days post injection (dpi) of CTX, and *eMyhc* and *Myh8* peaking at 5 dpi (Fig. 2F), reflecting sequential SC proliferation, differentiation and myofiber formation during muscle regeneration. Strikingly, the expression of *NRMLncR-v1/2* and *NRMLncR-v1* increased at 3 dpi and 5 dpi, although not significantly, while peaked at 7 dpi before gradually declining at 14 and 21 dpi. Notably, the expression of *NRMLncR-v1/2* and *NRMLncR-v1* was significantly increased by 40- and 80-fold respectively at 7 dpi compared with the uninjured muscle at 0 dpi (Fig. 2F). Consistently, a similar temporal induction pattern of *NRMLncR* was observed in an independent CTX- and glycerol-induced muscle injury model, with marked upregulation of *NRMLncR* at 7 dpi (GSE45577) (Fig. 2G and H). Together, these results demonstrate that *NRMLncR* expression dynamically correlated with the myogenic program.

### 3.3 | *NRMLncR* lacks protein-coding potential and is transcriptionally regulated by MRFs

Although the NCBI Gene Model annotated *NRMLncR* as a non-coding gene, the Ensembl Gene Model annotated it as a protein-coding gene predicted to encode an 88- and 56-amino acid micropeptides by *NRMLncR-v1* and *NRMLncR-v2,* respectively, indicating a conflict in BioType annotation. To assess the protein-coding potential of *NRMLncR-v1* and *NRMLncR-v2*, we performed Coding Potential Assessment Tool (CPAT) analysis and detected low coding probability scores for both isoforms, suggesting that both *NRMLncR-v1* and *NRMLncR-v2* were unlikely to be a protein-coding gene (Fig. 3A). To experimentally determine whether the two predicted open reading frames (ORF) of *NRMLncR-v1* and *NRMLncR-v2* produce detectable proteins, we cloned them into FLAG-tagged expression vectors with or without the predicted 5′ untranslated region (5′UTR), as well as constructs containing mutations in the ATG start codon and examined protein expression in HEK293 cells. While robust FLAG signals were detected from the positive control (pcDNA-Hey1-FLAG) by western blot analysis, no protein expression was observed from any of the ORF constructs of *NRMLncR-v1* and *NRMLncR-v2* (Fig. 3B). To exclude potential detection limitation of western blot for the predicted micropeptides, we further examined protein expression by immunofluorescence staining. Consistently, no FLAG signaling was detected in any ORF constructs of *NRMLncR-v1* and *NRMLncR-v2*, in contrast to the clear and robust FLAG signaling observed in the positive control (Fig. 3C and D). These results demonstrate that *NRMLncR* lacks protein-coding potential.

**Figure 3.**
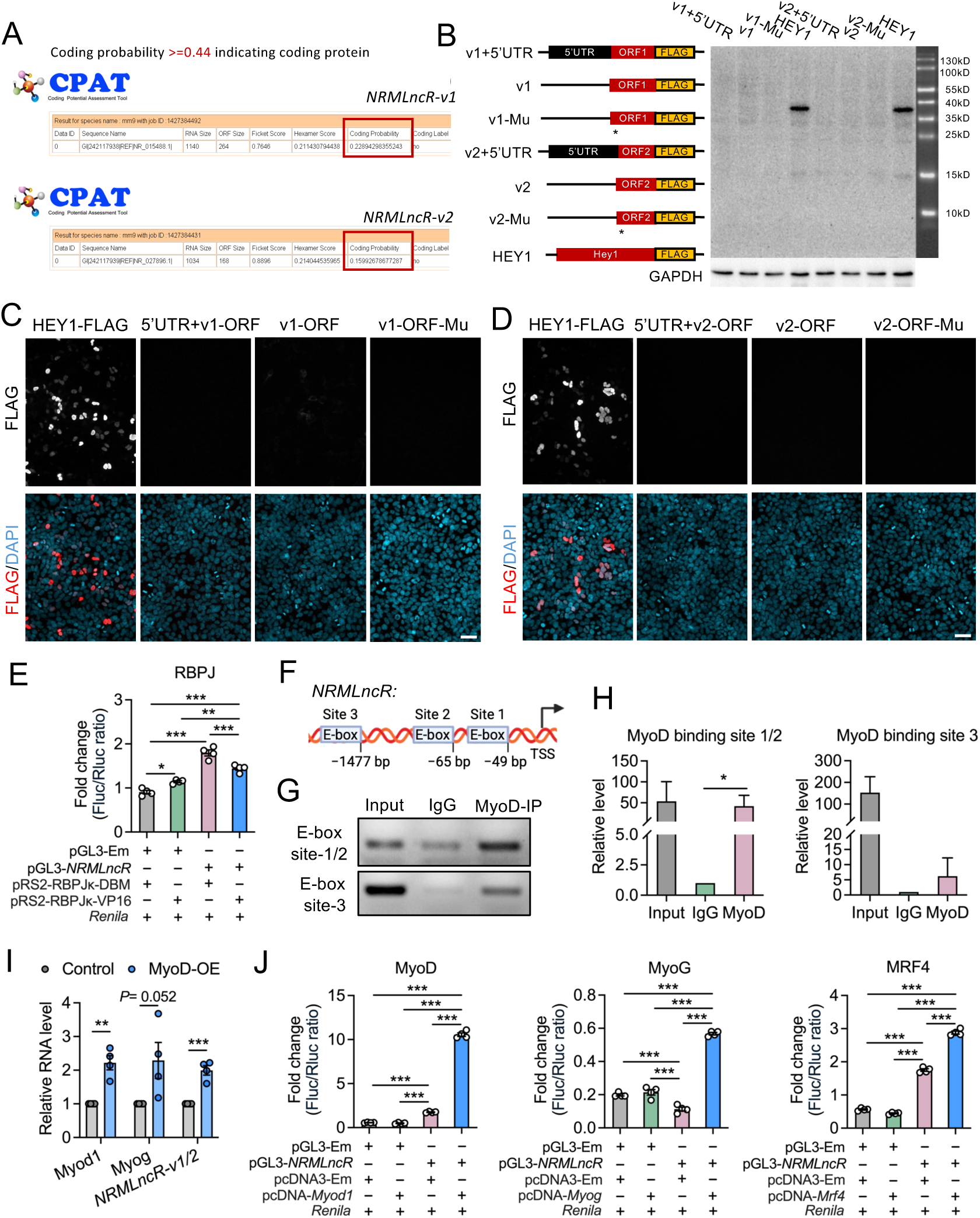
*NRMLncR* lacks protein-coding potential and is transcriptionally regulated by myogenic regulatory factors. (A) Coding Potential Assessment Tool (CPAT) analysis showing low protein-coding potential of *NRMLncR-v1* and *NRMLncR-v2*. Coding probability >=0.44 indicates coding protein. (B) Western blot analysis of FLAG tag to detect predicted proteins encoded by *NRMLncR-v1* and *NRMLncR-v2* open reading frames and their mutant ORF. FLAG-tagged HEY1 served as a positive control. (C and D) Immunofluorescence staining showing no detectable protein expression from the predicted *NRMLncR-v1* (C) and *NRMLncR-v2* (D) open reading frames in HEK293 cells, compared with a FLAG-tagged HEY1 positive control. Scale bar: 50 μm. (E) Luciferase reporter assays showing mildly inhibition of the *NRMLncR* promoter activity by RBPJ. n=4. (F) Schematic representation of putative E-box motifs within the *NRMLncR* promoter region. (G) ChIP-PCR and DNA gel analysis indicating the binding of MyoD to the E-box sites on the *NRMLncR* promoter. (H) ChIP-qPCR analysis showing enrichment of MyoD at the E-box regions of the *NRMLncR* promoter. n=3. (I) *NRMLncR-v1/2* expression levels in myoblasts overexpressing MyoD. n=4. (J) Luciferase reporter assays showing transcriptional activation of the *NRMLncR* promoter by MyoD, MyoG, and MRF4. n=4. Data represent mean ± SEM. One-way ANOVA multiple comparisons test (E, H, J), Student’s *t*-test (I): \**p* < .05; \*\**p* < .01; and \*\*\**p* < .001.

Given that the expression of *NRMLncR* was suppressed by Notch overexpression, we next examined whether *NRMLncR* transcription is directly regulated by RBPJ, a key transcription factor and primary mediator of the Notch signaling pathway. Notably, no canonical RBPJ binding site (5’-RGTGRGAA-3’) were identified in the promoter of *NRMLncR* (data not shown) and published datasets indicate that *NRMLncR* was not bound by RBPJ in ChIP-seq analyses.^60^ Dual-luciferase reporter assays showed that *NRMLncR* promoter activity was only mildly reduced (19.6%) in cells expressing constitutively active RBPJk-VP16 compared with cells expressing the mutant form RBPJk-DBM (Fig. 3E), suggesting that *NRMLncR* is not directly regulated by RBPJ. Notably, the *NRMLncR* promoter contains three putative E-box motifs (5’-CASCTG-3’) located at −49 bp, −65 bp and −1,477 bp (Fig. 3F), suggesting potential regulation by MRFs. To test this, we performed ChIP with MyoD antibody followed by PCR and DNA gel analysis, which revealed a marked enrichment of the promoter region containing the E-box sites bound by MyoD (Fig. 3G). Quantitative ChIP-qPCR analysis further demonstrated a 41.8- and 6.9-fold enrichment of MYOD binding at E-box site 1/2 and site 3 of *NRMLncR* promoter, respectively, compared to the IgG controls (Fig. 3H). Remarkably, overexpression of *Myod1* (MyoD-OE) in C2C12 myoblasts significantly upregulated the expression of *NRMLncR*-*v1/2* (Fig. 3I). Furthermore, Dual-luciferase reporter assays showed that expression of MyoD, MyoG, and MRF4 robustly activated the transcriptional activity of *NRMLncR* promoter (Fig. 3J). Together, these results indicate that *NRMLncR* is transcriptionally regulated by MRFs during myogenic differentiation.

### 3.4 | Knockdown of *NRMLncR* impairs myogenic differentiation

We next investigate the functional role of *NRMLncR* in myogenesis. To determine whether *NRMLncR* is required for myoblasts proliferation, we generated two C2C12 cell lines with stable expression of Lentivirual shRNAs targeting *NRMLncR-v1* or *NRMLncR-v1/2*, respectively, with a scramble shRNA as control (Fig. 4A). The efficiency of knockdown was validated by qPCR, which showed that both shRNAs significantly reduced the expression of total *NRMLncR-v1/2* and *NRMLncR-v1* in proliferating C2C12 myoblasts (Fig. 4B). Interestingly, *NRMLncR* knockdown led to decreased expression of key myogenic transcription factors, including *Myod1* and *Myog*, as well as the cell cycle regulator *Cyclin D1* (*Ccnd1*) (Fig. 4C). In contrast, the expression of the Notch target gene *Heyl* remained unchanged, whereas *β-Arrestin*, a gene involved in driving cell cycle exit and myogenic differentiation was reduced in *NRMLncR* knockdown C2C12 myoblasts (Fig. 4C). Despite these transcriptional changes, cell growth analysis revealed that myoblast number was not significantly affected by *NRMLncR* knockdown after 4 days of culture (Fig. 4D). These results indicate that *NRMLncR* silence alters myogenic gene expression but is not sufficient to disrupt myoblast proliferation.

**Figure 4.**
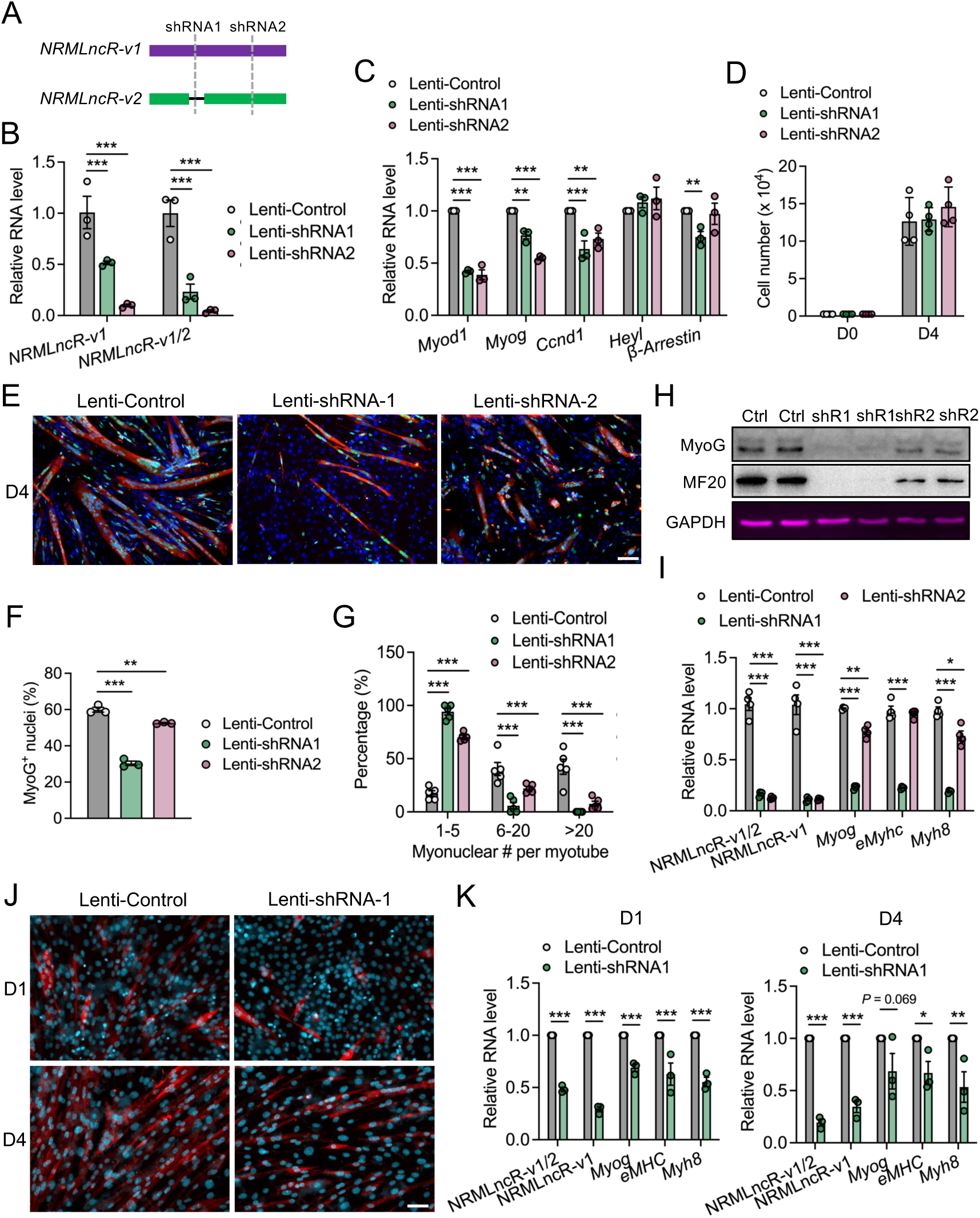
*NRMLncR* knockdown impairs myogenic differentiation. (A) Schematic illustration of two shRNA designs targeting *NRMLncR-v1* and total *NRMLncR-v1/2* for generating stable *NRMLncR* knockdown C2C12 cell lines. (B) Knockdown efficiency of *NRMLncR-v1* and *NRMLncR-v1/2* in stable *NRMLncR* knockdown C2C12 cell lines. n=3. (C) Relative mRNA expression levels in stable *NRMLncR* knockdown C2C12 cell lines. n=3. (D) Cell proliferation assay in control and *NRMLncR* knockdown C2C12 cell lines. n=4. (E) Immunofluorescence staining of MyoG and MF20 in control and *NRMLncR* knockdown C2C12 cultures after 4 days of differentiation. Scale bar: 50 μm. (F and G) Quantification of the percentage of MyoG^+^ nuclei (F) and myonuclei number in MF20^+^ myotubes (G) in control and *NRMLncR* knockdown C2C12 cultures after 4 days of differentiation. n=3 for F, n=5 for G. (H) Western blot analysis of myogenic proteins in control and *NRMLncR* knockdown C2C12 cultures after 4 days of differentiation. (I) Relative mRNA expression of myogenic genes in C2C12 cultures after 4 days of differentiation. n=4. (J) Immunofluorescence staining of MF20 in lentiviral-infected primary myoblasts after 1 and 4 days of differentiation. Scale bar: 50 μm. (K) Relative mRNA expression of myogenic genes in lentiviral-infected primary myoblasts after 1 and 4 days of differentiation. n=3. Data represent mean ± SEM. One- or Two-way ANOVA multiple comparisons test: \**p* < .05; \*\**p* < .01; and \*\*\**p* < .001.

We further examined the myogenic differentiation in C2C12 myoblasts with *NRMLncR* knockdown. Strikingly, upon induction of differentiation, *NRMLncR* knockdown cells formed fewer and smaller MF20^+^ myotubes compared to the control cells (Fig. 4E). Quantitative analysis revealed a substantial reduction in the proportion of MyoG^+^ nuclei (differentiation index) in Lenti-shRNA1 *NRMLncR* knockdown cells (60% in Lenti-Control versus 30% in Lenti-shRNA1), and a modest reduction in Lenti-shRNA2 *NRMLncR* knockdown cells (60% in Lenti-Control versus 52% in Lenti-shRNA2) (Fig. 4F). Moreover, the myogenic fusion index was markedly decreased in both *NRMLncR* knockdown lines after 4 days of differentiation, as indicated by a 5.5- and 4.1-fold increase in the proportion of myotubes containing 1-5 myonuclei in Lenti-shRNA1 and Lenti-shRNA2 *NRMLncR* cells, respectively (Fig. 4G). In contrast, myotubes containing more than 20 myonuclei were rarely observed in either *NRMLncR* knockdown group (Fig. 4G). At the molecular level, western blot analysis showed markedly reduced MyoG and MF20 protein levels following *NRMLncR* knockdown (Fig. 4H), and qPCR analysis confirmed decreased expression of myogenic markers *Myog*, *eMyhc*, and *Myh8* (Fig. 4I). To validate these findings in a more physiological context, we generated SC-derived primary myoblasts with *NRMLncR-v1* knockdown. Time-course analysis demonstrated that *NRMLncR-v1* knockdown impaired myogenic differentiation of primary myoblasts at both early (day 1) and later (day 4) stages, with much less MF20^+^ myotubes formed (Fig. 4J). Consistently, the expression of myogenic markers *Myog*, *eMyhc*, and *Myh8* was significantly reduced in *NRMLncR-v1* knockdown primary myoblasts at both 1 and 4 days of differentiation (Fig. 4K). Together, these results demonstrate that *NRMLncR* is required for efficient myogenic differentiation.

### 3.5 | *NRMLncR* overexpression enhances myogenic differentiation and muscle growth

To determine whether *NRMLncR* overexpression promotes myogenic differentiation, we generated an adenovirus expressing *NRMLncR-v1* (Ad-NRMLncR-v1) and infected SC-derived primary myoblasts. Efficient overexpression of *NRMLncR-v1* in primary myoblasts was confirmed by qPCR (Fig. 5A). After 4 days of differentiation, *NRMLncR-v1*-overexpressing cells formed larger and more multinucleated MF20^+^ myotubes compared with control cells infected with Ad-GFP (Fig. 5B). Specifically, the percentage of myotubes containing 1-5 myonuclei decreased by 28.3% (78.7% in Ad-GFP versus 56.5% in Ad-NRMLncR-v1), whereas the percentage of myotubes containing more than 20 myonulcei increased 3.1-fold in *NRMLncR-v1*-overexpressing cells (5.7% in Ad-GFP versus 17.9% in Ad-NRMLncR-v1) (Fig. 5C). To assess the effect of *NRMLncR* overexpression on myogenesis *in vivo*, adenovirus expressing *NRMLncR-v1* or GFP was injected into the TA muscles of 3-week-old mice (Fig. 5D). At 14 days post injection (dpi), TA muscle weight showed a modest increase in Ad-NRMLncR-v1-infected muscles compared to the Ad-GFP-infected controls (Fig. 5E). Histological analysis of TA muscle cross-sections revealed that *NRMLncR-v1* overexpressing myofibers were morphologically larger than those in control muscles (Fig. 5F). Quantitative analysis confirmed an increase in myofiber cross-sectional area (CSA) in Ad-NRMLncR-v1-infected muscles at 14 dpi (Fig. 5G).

**Figure 5.**
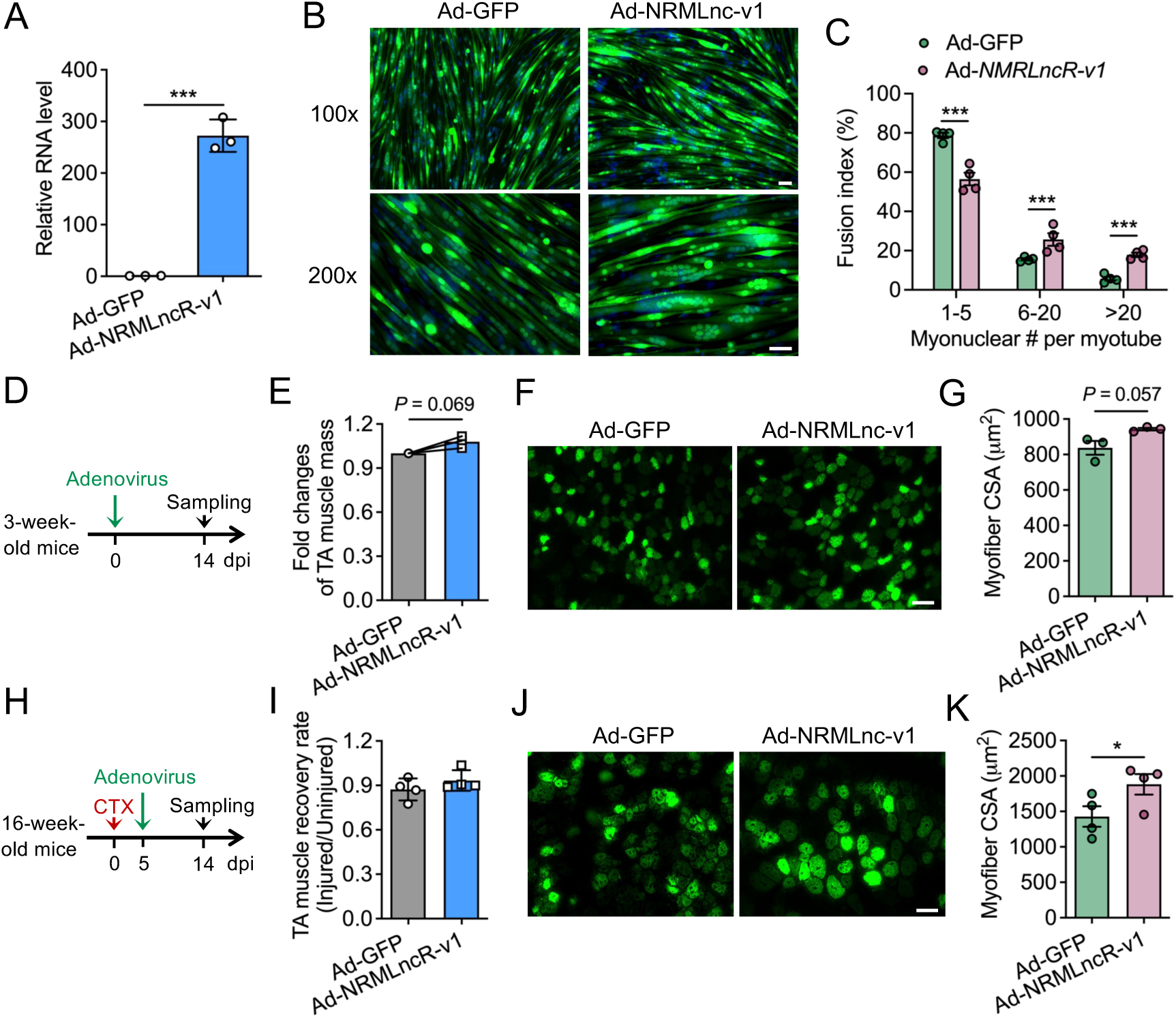
*NRMLncR* overexpression enhances myogenic differentiation and muscle growth. (A) Overexpression efficiency of *NRMLncR-v1* in primary myoblasts following adenoviral infection. n=3. (B) Immunofluorescence staining of GFP in control (Ad-GFP) and *NRMLncR-v1*-overexpressing (Ad-NRMLncR-v1) myotubes after 4 days of differentiation. Scale bar: 50 μm. (C) Quantification of the percentage of myonuclei number per myotube in control and *NRMLncR-v1*-overexpressing myotubes after 4 days of differentiation. n=4. (D) Schematic illustration of the experimental design for *NRMLncR-v1* overexpression in the tibialis anterior (TA) muscle of 3-week-old mice. (E) TA muscle weight of 3-week-old mice receiving Ad-GFP and Ad-NRMLncR-v1 at 14 days post injection (dpi). n=3. (F) Immunofluorescence staining of GFP on TA muscle cross-sections of 3-week-old mice receiving Ad-GFP or Ad-NRMLncR-v1 at 14 dpi. Scale bar: 50 μm. (G) Quantification of myofiber cross-sectional area (CSA) in TA muscles from 3-week-old mice 14 days after Ad-GFP or Ad-NRMLncR-v1 injection. n=3. (H) Schematic illustration of the experimental design for *NRMLncR-v1* overexpression in TA muscle of 16-week-old mice following CTX-induced injury. (I) TA muscle weight of mice receiving Ad-GFP or Ad-NRMLncR-v1 and CTX injury at 14 dpi. n=4. (J) Immunofluorescence staining of GFP on TA muscle cross-sections of mice receiving Ad-GFP or Ad-NRMLncR-v1 and CTX injury at 14 dpi. Scale bar: 50 μm. (K) Quantification of myofiber CSA from mice receiving Ad-GFP or Ad-NRMLncR-v1 and CTX injury at 14 dpi. n=4. Data represent mean ± SEM. Student’s *t*-test: \**p* < .05; and \*\*\**p* < .001.

To further evaluate the role of *NRMLncR* during muscle regeneration, we employed a CTX-induced muscle injury model following with adenoviral injection. Although the TA muscle recovery rate, calculated as the ratio of injured to contralateral uninjured muscle weight, was not significantly altered by *NRMLncR* overexpression (Fig. 5H and I), histological analysis revealed enlarged GFP-positive myofibers in Ad-NRMLncR-v1-infected muscles at 14 dpi (Fig. 5J). Consistently, quantitative analysis demonstrated a significantly increase in myofiber CSA in Ad-NRMLncR-v1-infected muscles compared to the Ad-GFP controls (Fig. 5K). Together, these *in vitro* and *in vivo* findings indicate that *NRMLncR* overexpression enhances myogenic differentiation and promotes muscle growth.

### 3.6 | *NRMLncR* regulates neighboring gene expression and interacts with RNA-binding protein CELF1

We next sought to explore the molecular mechanism underlying *NRMLncR* function during myogenic differentiation. Because the subcellular localization of lncRNAs is often linked to their function, we first examined the localization of *NRMLncR*. qPCR analysis of the fractionated RNA extracted from differentiated C2C12 cells showed that non-coding RNA *U6* was predominantly enriched in the nuclear fraction, whereas *18s* rRNA and *β-actin* mRNA were present at comparable levels in both cytoplasmic and nuclear fractions. Strikingly, *NRMLncR-v1* and total *NRMLncR-v1/2* were predominantly localized in the cytoplasmic fraction of C2C12 cells, although detectable levels were also present in the nucleus (Fig. 6A).

**Figure 6.**
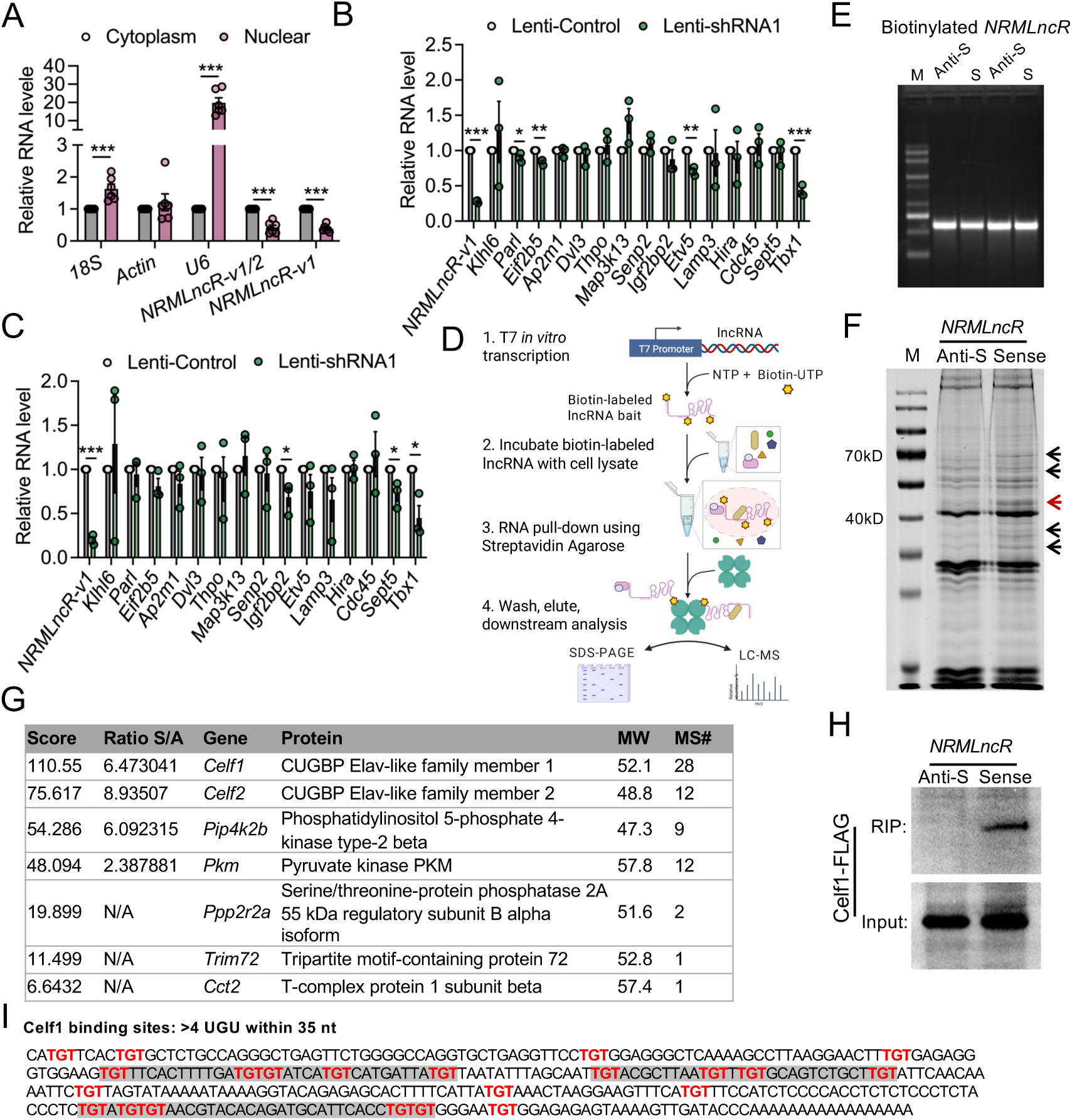
*NRMLncR* regulates neighboring gene expression and interacts with CELF1. (A) qPCR analysis showing subcellular localization of *NRMLncR-v1* and *NRMLncR-v1/2* in the cytoplasmic and nuclear fractions of C2C12 cells. n=6. (B and C) Relative mRNA expression of neighboring genes of the *NRMLncR* locus in lentiviral-infected primary myoblasts after 1 (B) and 4 (C) days of differentiation. n=3. (D) Schematic illustration of the RNA pull-down and mass spectrometry workflow used to identify *NRMLncR*-interacting proteins. (E) Electrophoresis analysis showing *in vitro*–transcribed, biotin-labeled sense and antisense *NRMLnc* RNAs. (F) Coomassie blue staining of SDS-PAGE gel showing proteins precipitated by biotinylated sense and antisense *NRMLncR* RNAs in differentiated C2C12 myotubes. n=3 independent experiments. Black arrows indicates protein bands enriched in sense *NRMLncR* RNA pull down. Red arrow indicate the protein band for MS analysis. (G) Top candidate *NRMLncR*-interacting proteins identified by mass spectrometry. (H) RNA pull-down followed by immunoblotting validating the interaction between *NRMLncR* and CELF1. n=3 independent experiments. (I) Predicted UGU-rich elements (GREs) within *NRMLncR* that mediate interaction with CELF1. Data represent mean ± SEM. Student’s *t*-test: \**p* < .05; and \*\*\**p* < .001.

Nuclear LncRNAs often function *in cis* acting as enhancer-like elements to regulate the expression of their neighboring genes on the same chromosome.^61^ To determine whether *NRMLncR* regulates the expression of neighboring genes *in cis*, we examined the expression of fifteen genes located on both sides of the *NRMLncR* locus (−1.2M to +2.5M) in *NRMLncR-v1* knockdown C2C12 myoblasts. Notably, the expression of *Tbx1,* a gene with an essential role in myogenesis, was markedly reduced in proliferating C2C12 cells upon *NRMLncR-v1* knockdown (Fig. 6B). In addition, the expression of *Etv5*, *Eif2b5*, and *Parl* was significantly decreased, although the magnitude of reduction was comparatively modest. At 4 days of differentiation, most neighboring genes were not significantly affected by *NRMLncR-v1* knockdown. In contrast, the expression of *Tbx1* remained persistently downregulated in differentiated C2C12 myotubes, while *Igfbp2* and *Sept5* exhibited mild decreases in expression (Fig. 6C). These results suggest that *NRMLncR* might influence myogenesis by modulating the transcriptional regulation of nearby genes.

To determine whether *NRMLncR* functions through interacting with other proteins, we performed RNA pull-down assays using biotinylated sense and antisense *NRMLncR* RNA synthesized *in vitro,* followed by mass spectrometry (MS) analysis (Fig. 6D and E). Coomassie blue staining of SDS-PAGE gel revealed several protein bands enriched in the sense *NRMLncR* RNA pull-down from C2C12 myotubes, which are distinguishable from the antisense control (Fig. 6F). We focused on a prominent protein band with high abundance located between 40 and 55 kDa. Further MS analysis of this band identified several candidate *NRMLncR*-interacting proteins, among which CELF1 (CUGBP1) showed highest enrichment, with the CELF2 (CUGBP2) ranking second (Fig. 6G). To validate the interaction between *NRMLncR* and CELF1, RNA pull-down assays was performed using HEK293T cells expressing CELF1, followed by immunoblotting analysis. Notably, a strong CELF1 signal was detected in the eluates from the sense *NRMLncR* RNA pull-down, whereas no detectable signal was observed in the antisense RNA control (Fig. 6H), suggesting a direct interaction between *NRMLncR* and CELF1. In addition, bioinformatic analysis identified multiple UGU-rich elements (GREs) within *NRMLncR*, suggested putative binding sites for CELF1 interaction (Fig. 6I).^62^ Together, these results indicate that *NRMLncR* physically interacts with the RNA-binding protein CELF1.

## 4 | DISCUSSION

In this study, we identified *NRMLncR* (*A930003A15Rik*) as a previously uncharacterized myocyte-enriched lncRNA whose expression is suppressed by Notch activation yet robustly induced during SC activation and differentiation and injured-induced regeneration. Functionally, *NRMLncR* is required for efficient myogenesis, evident by that *NRMLncR* knockdown markedly reduces myogenic gene expression and impairs myotube formation and fusion, whereas its overexpression promotes myogenic differentiation *in vitro* and results in increased myofiber size *in vivo*. Mechanistically, the transcription of *NRMLncR* is directly regulated by core myogenic program as MRFs occupy and activate its promoter. *NRMLncR* is predominantly cytoplasmic yet it is also detectable in nucleus and interacts with RNA-binding protein CELF1. These findings position *NRMLncR* as a new component linking canonical Notch-MRF signaling to lncRNA-mediated regulation of myogenic progression.

Our observations indicate that *NRMLncR* primarily impacts differentiation and fusion rather than myoblast proliferation, consistent with its induction during the differentiation program. Although *NRMLncR* knockdown reduced expression of key myogenic regulators MyoD and MyoG and altered select cell-cycle genes, total cell number over several days in growth conditions was not significantly changed, indicating that *NRMLncR* is not essential for maintaining the proliferative capacity in myoblasts under the present assay conditions. A plausible explanation is that *NRMLncR* functions most strongly at the transition into differentiation, such as cell-cycle exit, myogenic lineage specialization, and fusion competence. In this scenario, small changes in regulatory networks induced by *NRMLncR* knockdown can produce outsized effects on MyoG induction and myotube formation, even if proliferation remains buffered by redundant pathways. This interpretation is supported by the expression dynamics of *NRMLncR* during myogenic differentiation and regeneration and by the pronounced defects in differentiation and fusion upon knockdown, alongside improved fusion with overexpression. An important related question is whether *NRMLncR* participates earlier during SC activation from quiescence. Our data are consistent with a role in myogenic progression yet do not resolve whether *NRMLncR* is required for quiescence exit or late-stage myotube maturation. It will be worth of addressing this utilizing *in vivo* or *ex vivo* SC-specific perturbation approach, paired with cell-cycle kinetics and cell state transitions during regeneration.

Although we identified *NRMLncR* in Notch-activated myoblast, upstream regulation studies suggest *NRMLncR* is controlled primarily by MRFs, while Notch likely represses *NRMLncR* indirectly through its established anti-differentiation circuitry.^17, 20^ *NRMLncR* is among the most downregulated lncRNAs upon Notch activation, and pharmacologic Notch activation markedly suppresses *NRMLncR* expression, supporting the idea that *NRMLncR* is Notch-responsive. However, *NRMLncR* does not appear to be a canonical RBPJ direct target as no classic RBPJ motif was identified in the promoter and activated RBPJ produced only a modest repression in reporter assays. In contrast, the *NRMLncR* promoter contains E-box motifs bound by MyoD, and MyoD/MyoG/MRF4 robustly activate *NRMLncR* promoter activity, indicating direct control by the myogenic transcriptional cascade. Given the well-established role of Notch signaling in maintaining SC quiescence and inhibiting differentiation,^18, 19, 25^ an attractive model is that Notch suppresses *NRMLncR* primarily by limiting MRF activity and/or via downstream transcriptional repressors (for example: HES/HEY family factors) that antagonize the myogenic program.^36^ Whether HES/HEY factors or other Notch-induced transcriptional regulators directly influence *NRMLncR* transcription remains untested and warrants future investigation, particularly in primary SCs across defined activation and differentiation stages.

In this study, we provide evidence that *NRMLncR* may exert regulatory functions through multiple mechanisms during myogenesis. We found that *NRMLncR* localizes to both the cytoplasm and the nucleus, suggesting the potential to engage in distinct layers of gene regulation.^61^ Notably, knockdown of *NRMLncR* altered the expression of several neighboring genes, with *Tbx1* emerging as the most consistently and robustly downregulated target, supporting a possible *cis*-regulatory role for *NRMLncR*. This observation is particularly intriguing given the established importance of Tbx1 in myogenic regulation and muscle development.^63^ Previous studies have shown that muscle-enriched lncRNAs such as *linc-MD1* and *lnc-mg* regulate the timing of muscle differentiation primarily by functioning as competing endogenous RNAs to fine-tune pro-differentiation gene expression programs.^39,40^ In contrast, regulation of neighboring gene transcription by lncRNAs during myogenesis has not been well documented.^37, 38^ Thus, our findings identify a previously unappreciated mode of lncRNA-mediated regulation in muscle cells and suggest that *NRMLncR* may contribute to myogenic control, at least in part, through modulation of local gene expression. Nonetheless, it remains to be determined whether Tbx1 serves as a direct downstream effector of *NRMLncR* in regulating myogenesis. Future studies will be required to dissect the molecular mechanisms underlying *NRMLncR*-mediated *cis* regulation, including its chromatin interactions and regulatory complexes, and to establish the functional contribution of Tbx1 to *NRMLncR*-dependent myogenic outcomes.

Moreover, unbiased RNA pull-down coupled with LC-MS identified CELF1 and CELF2 as enriched *NRMLncR*-interacting proteins, and follow-up validation supports a direct *NRMLncR*-CELF1 interaction. CELF1 is a well-studied regulator of RNA fate through alternative splicing, mRNA stability/decay, and translation.^64–66^ Dysregulation of CELF1 has been implicated in myotonic dystrophy, consistent with broad roles in inhibiting myogenic RNA programs.^67^ Thus, we propose that *NRMLncR* may act as an RNA “modulator” of CELF1, either sequestering CELF1 away from pro-differentiation transcripts, altering CELF1 binding specificity and affinity, or reshaping CELF1-containing ribonucleoprotein complexes,^68^ thereby protecting myogenic gene expression during the differentiation process. A key limitation is that we have not yet defined the functional consequence of this interaction. Future studies should be considered to investigate the changes in CELF1 occupancy on specific mRNAs during myogenic differentiation, the effects on transcript stability, splicing, and translation, or whether CELF1 also regulates *NRMLncR* turnover. In addition, the presence of other enriched bands in the pull-down suggests that *NRMLncR* may interact with additional proteins and underlie mechanisms beyond CELF1, which should be mapped by orthogonal approaches.

In summary, *NRMLncR* emerges as a myogenic lncRNA positioned at the Notch-MRF interface to facilitate differentiation and regeneration, but its conservation and translational potential remain open questions. Integrating our findings, we propose a working model in which declining Notch signaling during activation/differentiation permits MRF-driven induction of *NRMLncR*; *NRMLncR* then promotes differentiation and fusion, at least in part, through cytoplasmic interaction with CELF1 and potentially other RNA-binding proteins. Of note, whether *NRMLncR* is conserved in humans is unclear. Our preliminary sequence-based searches did not identify an obvious human ortholog, raising the possibility that *NRMLncR* may be rodent-specific or that functional conservation occurs at the level of genomic context, secondary structure, or pathway logic rather than primary sequence. Future work should therefore determine the conservation by synteny or structure and assess expression in human myoblast differentiation, establish *in vivo* necessity using loss-of-function models during regeneration, and define the mechanistic outputs of the *NRMLncR*-Tbx1 and *NRMLncR*-CELF1 axis on RNA transcription and processing to clarify how *NRMLncR* tunes myogenic gene expression programs.

## Supporting information

Supplemental Datasheet 1

## DATA AVAILABILITY STATEMENT

All data reported in this manuscript will be shared by the lead contact upon request. No original code was reported that needed to reanalyze the data generated by this study. The microarray data generated in this study have been provided as supplemental datasheet. Any additional information required to reanalyze the data reported in this manuscript is available from the corresponding author upon request.

## AUTHOR CONTRIBUTIONS

**Yufen Li**: Formal analysis; Visualization; Writing—original draft. **Yumei Zhou**: Formal analysis; Visualization; Writing—review and editing. **Qing Ying Li**: Formal analysis; Visualization; Writing—review and editing. **Justine Arrington**: Data curation; Formal analysis; Methodology. **Mostafa F. Abdelhai**: Formal analysis; Visualization. **Yubo Wang**: Formal analysis; Validation. **Junxiao Ren**: Writing—review and editing. **Yung-Yu Cheng**: Writing—review and editing. **Mingyi Xie**: Resources; Writing—review and editing. **W. Andy Tao**: Resources; Methodology; Writing—review and editing. **Shihuan Kuang**: Resources; Supervision; Funding acquisition; Writing—review and editing. **Feng Yue**: Conceptualization; Data curation; Software; Formal analysis; Supervision; Funding acquisition; Project administration; Validation; Investigation; Visualization; Methodology; Writing—review and editing.

## ACKNOWLEDGMENTS

This work was supported by grants from the National Institutes of Health NIH-R01DK136722 to FY, NIH-R01DK132819 and NIH-R01AR060652 to SK, NIH-R35GM128753 to MX, the Muscular Dystrophy Association MDA516161 to FY, and the University of Florida Research Startups to FY. We thank Dr. Jennifer Freeman and Sara Wirbisky at Purdue University for helping with microarray analysis, Dr. Yefei Wen for generating the primary myoblasts from *Rosa-NICD* mcie and assisting microarray analysis, and Dr. Yong-Xu Wang at the University of Massachusetts Medical School for providing the Ad-Cre and Ad-GFP adenovirus. We are grateful to Purdue Flow Cytometry and Cell Separation Facility for assistance on flow cytometry analysis.

