## Supplemental Table 1 for "*NRMLncR,* a myocyte-enriched long non-coding RNA, enhances myogenesis in mouse"

### Supplemental Material

#### Supplementary Table 1

**List of primers used in this study**

| Application | Gene name | Primer sequence (5'-3') |
|---|---|---|
| qPCR | *18s* | F: AGTCCCTGCCCTTTGTACACA |
| | | R: CGATCCGAGGGCCTCACTA |
| qPCR | *U6* | F: CTCGCTTCGGCAGCACA |
| | | R: AACGCTTCACGAATTTGCGT |
| qPCR | *β-actin* | F: GGCTGTATTCCCCTCCATCG |
| | | R: CCAGTTGGTAACAATGCCATGT |
| qPCR | *NRMLncR-v1/2* | F: CGGATAGCAGAGGCAGAGATGAG |
| | | R: CAGCCCAGGAGGAGACTTCAT |
| qPCR | *NRMLncR-v1* | F: AAGATGGTGACCTTGCGTCCT |
| | | R: GCATCAGGCGTGTGTCCAG |
| qPCR | *Pax7* | F: CGACTCTGGATTCGTCTCC |
| | | R: GGCCTTGGCCAAGAGGG |
| qPCR | *MyoD* | F: GGCTACGACACCGCCTACTA |
| | | R: CGACTCTGGTGGTGCATCTG |
| qPCR | *MyoG* | F: TGCCCAGTGAATGCAACTCC |
| | | R: TTGGGCATGGTTTCGTCTGG |
| qPCR | *eMyhc* | F: AAAAGGCCATCACTGACGC |
| | | R: CAGCTCTCTGATCCGTGTCTC |
| qPCR | *Myh8* | F: GGAGAGGATTGAGGCCCAAAA |
| | | R: CACGGTCACTTTCCCTCCATC |
| qPCR | *Hey1* | F: TGAATCCAGATGACCAGCTACTGT |
| | | R: TACTTTCAGACTCCGATCGCTTAC |
| qPCR | *Hey2* | F: AAGCGCCCTTGTGAGGAAAC |
| | | R: GGTAGTTGTCGGTGAATTGGAC |
| qPCR | *Hes2* | F: ACAATTACCCTGGGCACGCTAC |
| | | R: CCTGTAGCCTGGAGCATCTTCAAA |
| qPCR | *Heyl* | F: CAGATGCAAGCCCGGAAGAA |
| | | R: ACCAGAGGCATGGAGCATCT |
| qPCR | *Follistatin* | F: GCCAGTGACAATGCCACATACG |
| | | R: CTTCCTCCGTTTCTTCCGAGATG |
| qPCR | *Ap2m1* | F: GACATCGGGAGGAATGCTGTG |
| | | R: TGGACCGCTTAACATGGAAGA |
| qPCR | *Cdc45* | F: GATTTCCGCAAGGAGTTCTACG |
| | | R: TACTGGACGTGGTCACACTGA |
| qPCR | *Dvl3* | F: GTCACCTTGGCGGACTTTAAG |
| | | R: AAGCAGGGTAGCTTGGCATTG |
| qPCR | *Eif2b5* | F: AGTTCTAGTGGCCGATAGCTT |
| | | R: AGCAGCAAAAGACAAATGTTTCC |
| qPCR | *Etv5* | F: TCAGTCTGATAACTTGGTGCTTC |
| | | R: GGCTTCCTATCGTAGGCACAA |
| qPCR | *Hira* | F: CCACCGTTCGGGGGATAAG |
| | | R: GGCAACACATACCACATCACAG |
| qPCR | *Igf2bp2* | F: GTCCTACTCAAGTCCGGCTAC |
| | | R: CATATTCAGCCAACAGCCCAT |
| qPCR | *Klhl6* | F: GCTTGGAAGGACCCTTAGCAC |
| | | R: CGTCTGTCAAAGCATTTTCCTCT |
| qPCR | *Lamp3* | F: CAAGGACAGATCAACGACCTC |

| | | |
|---|---|---|
| | | R: GCCTGCTTCCATTTAGGACTTC |
| qPCR | *Map3k13* | F: CCCGACCTCATCTCCACAG |
| | | R: TGGAAACAGGGATCATAGGGTT |
| qPCR | *Parl* | F: TACGGCCACAAAAGGAAGGAA |
| | | R: TTCGCAGCTATGATGCCTGTC |
| qPCR | *Senp2* | F: GCTGGCTAAGGTTCTCGGC |
| | | R: CTGGGATCTCATCAGTGTCCA |
| qPCR | *Sept5* | F: GAAAGGTTTCGACTTCACGCT |
| | | R: CCGGTCCTTATACAGGTCGGT |
| qPCR | *Thpo* | F: GGCCATGCTTCTTGCAGTG |
| | | R: AGTCGGCTGTGAAGGAGGT |
| qPCR | *Ccnd1* | F: GCGTACCCTGACACCAATCTC |
| | | R: CTCCTCTTCGCACTTCTGCTC |
| qPCR | *β-arrestin1* | F: CCTGCATCAGCCAGATGAAG |
| | | R: GGTGTTTCAGGATGGCCTTC |
| PCR | *NRMLncR* | F: ACCTTGAGTGATCTTCAGGGAGA |
| | | R: GGCATCAGGCGTGTGTCCAG |
| 5'UTR+ORF | *NRMLncR* | F: CGCTCTAGAAGATGGTAAATTCCCTTAGCA |
| | | R: TTTGGATCCTCGATGGTTTGCAGACTGCAGC |
| ORF | *NRMLncR-v1* | F: CGCTCTAGAATGGAGGCCAGAGAAGATG |
| | *NRMLncR-v1-mutant* | F: CGCTCTAGATAGGAGGCCAGAGAAGATG |
| ORF | *NRMLncR-v2* | F: TATTCTAGAATGGTGGTGCGGATAGCAG |
| | *NRMLncR-v2-mutant* | F: TATTCTAGATAGGTGGTGCGGATAGCAG |
| ORF | *Hey1* | F: CGCTCTAGAATGAAGAGAGCTCACCCAGACT |
| | | R: TTTGGATCCGAAAGCTCCGATCTCTGTCC |
| Promoter | *NRMLncR* | F: TTTACGCGTAGCTTTGAATCAGCAGGACTGTC |
| | | R: GGGCTCGAGTAAGGGAATTTACCATCTAAGG |
| ChIP-PCR | E-box site 1/2 | F: CAGCAAGACAGAGCATTCTTCAC |
| | | R: CAAACGGTAAACGGCTCAAAAG |
| ChIP-PCR | E-box site 3 | F: AGAACCCTGGATCTAGTAGCTGAG |
| | | R: GGCTCTGAGTTGACCTCTTCATT |
| shRNA | NRMLncR-shRNA-1 | F:CCGGGGAGGTTTAACTATACCAATGCCACACCCATTGGTATAGTTAAACCTCCTTTTTG |
| | | R:AATTCAAAAAGGAGGTTTAACTATACCAATGGGTGTGGCATTGGTATAGTTAAACCTCC |
| shRNA | NRMLncR-shRNA-2 | F:CGGGGAGGTTTAACTATACCAATGCCACACCCATTGGTATAGTTAAACCTCCTTTTTG |
| | | R:TCGAGAAAAAGCAGTCTGCTTGTATTCAACACTCGAGTGTTGAATACAAGCAGACTGCT |
| In vitro transcription | *NRMLncR-v1* | F: TAATACGACTCACTATAGGGAGATGGTAAATTCCCTTAGCA |
| | | R: GGGTATCAACTTTTACTCTCTCC |
| Adenoviral OE | *NRMLncR-v1* | F: TTTGCGGCCGCAGATGGTAAATTCCCTTAGCA |
| | | R: GCGGATATCGGGTATCAACTTTTACTCTCTCC |
